# Topography-dependent gene expression and function of common cell archetypes in large and small porcine airways

**DOI:** 10.1101/2021.03.16.435690

**Authors:** Alejandro A. Pezzulo, Andrew L. Thurman, Xiaopeng Li, Raul Villacreses, Wenjie Yu, Steven E. Mather, Guillermo S. Romano-Ibarra, David K. Meyerholz, David A. Stoltz, Michael J. Welsh, Ian M. Thornell, Joseph Zabner

## Abstract

The small airways of humans are affected early in several lung diseases. However, because they are relatively inaccessible, little is known about the epithelial cells that line these airways. We performed a single cell RNA-seq census of small and large airways of wild-type pigs and pigs with disrupted cystic fibrosis transmembrane conductance regulator (*CFTR*) gene. The sequencing data showed that small airway epithelia had similar major cell types as large airways but no ionocytes; moreover, lack of *CFTR* expression had minimal effect on the transcriptome. Small airway epithelial cells expressed a different transcriptome than large airway cells. Quantitative immunohistochemistry showed that small airway basal cells participate in epithelial barrier function. Finally, sequencing data and in vitro electrophysiologic studies suggest that small airway epithelia have a water and ion transport advantage. Our data highlight the archetypal nature of basal, secretory, and ciliated airway cells with location_-_dependent gene expression and function.

## Introduction

The airways of humans and other large mammals can be classified into two broad types: 1) “Large” or proximal airways including the trachea and the lobar, segmental, and sub-segmental bronchi, which contain submucosal glands and cartilage, and 2) “Small” or distal airways that include the non-respiratory and respiratory bronchioles. The small airways are devoid of submucosal glands and cartilage, and are < 2 mm diameter in humans (Brown et al., 1969; Carr et al., 2017; Hogg et al., 1968; Hogg et al., 2017; Hyde et al., 2009; Macklem and Mead, 1967a, b). The early events in the pathogenesis of many respiratory diseases including asthma, chronic obstructive pulmonary disease (COPD), and cystic fibrosis (CF) may occur in the small airways (Ratjen, 2012; Shaw et al., 2002; Tiddens et al., 2010; Zinellu et al., 2019)

The cellular composition and structure of large airway epithelia have been well_-_established through histological and immunofluorescence imaging, flow cytometry, single-cell RNA sequencing, and other techniques (Bonser et al., 2021; Hackett et al., 2008; Jackson et al., 2020; Maestre-Batlle et al., 2017; Montoro et al., 2018; Nakajima et al., 1998; Okuda et al., 2020; Plasschaert et al., 2018). The large airway epithelium is a pseudostratified columnar epithelium consisting of basal cells in the basal layer, and secretory and ciliated cells residing in an apical/surface layer. Basal cells function as progenitors of other epithelial cells (Boers et al., 1998; Hong et al., 2004a, b; Lynch et al., 2019; Rock et al., 2009) and regulate adhesion of the epithelium to the basement membrane (Evans and Plopper, 1988; Hawkins et al., 2021; Nakajima et al., 1998). Secretory cells can be further divided into two subtypes, club cells and goblet cells. Club cells (previously called Clara cells) (Winkelmann and Noack, 2010) are non-ciliated exocrine cells with membrane-bound secretory granules containing proteins important for airway host defense and regulation of inflammation, ion transport, and airway surface liquid (ASL) pH (Chinet et al., 1997; Snyder et al., 2010; Van Scott et al., 1987; Zuo et al., 2018). Goblet cells are specialized secretory cells containing mucin granules that contribute to the protective layer of the airway surface upon secretion (Atherton et al., 2003; Boucherat et al., 2013; Boucherat et al., 2012; Chen et al., 2009; Danahay et al., 2015; Hayashi et al., 2004; Ma et al., 2017; Pezzulo et al., 2019). Ciliated cells contain hair-like structures, called cilia, that extend into the airway lumen and work in concert with the airway surface fluid to clear the airway of foreign particles and pathogens (Brody, 2004; Horani et al., 2018; Nanjundappa et al., 2019; Whitsett, 2018; You et al., 2004). In addition to the three major cell types, airway epithelia also contain rare cell types, including pulmonary neuroendocrine cells (PNECs), tuft or brush cells, and CFTR_-_rich ionocytes (Bankova et al., 2018; Krasteva et al., 2012; Montoro et al., 2018; Ouadah et al., 2019; Perniss et al., 2020; Perniss et al., 2021; Plasschaert et al., 2018; Scudieri et al., 2020; Sui et al., 2018).

Whereas there is abundant literature describing the histopathology of airway disease in the small airways, less is known about the origins, function, and role in disease of small airway epithelia and their cellular components (Bhowmick and Gappa-Fahlenkamp, 2016; Hackett et al., 2012; O’Beirne et al., 2018; Okuda et al., 2019; Shamsuddin and Quinton, 2012, 2014; Tilley et al., 2011; Vucic et al., 2014; Yang et al., 2017; Zuo et al., 2020; Zuo et al., 2018). The analysis of human small airways has been limited by their poor accessibility to measurements or sampling in vivo in living subjects. Changes in airway resistance caused by the large airways are easily and reproducibly measured with pulmonary function testing; in contrast, small airways have a minimal and poorly measurable contribution to overall airflow resistance (Brown et al., 1969; Carr et al., 2017; Hogg et al., 1968; Hogg et al., 2017; Macklem and Mead, 1967b). Moreover, whereas sampling of large airway epithelia via bronchoscopic forceps or brush biopsy is straightforward, accurate sampling of 2 mm diameter and smaller airways (8-10^th^ or further generations of the airway tree) is challenging (Wang et al., 2019).

Other limitations to the study of small airway epithelia include the marked inter-species differences between large mammals and small animal models such as mice. Whereas mice have small-diameter airways morphologically similar to human small airways, they are functionally different, for example: 1) mice without CFTR do not develop lung disease resembling cystic fibrosis (reviewed in (Grubb and Boucher, 1999; Guilbault et al., 2007)), 2) mice express different ion transporters important for airway disease than humans (e.g., the acid transporter ATP12A (Chen et al., 2010; Shah et al., 2016), and 3) mouse airway epithelial cells may follow different developmental lineages than those of humans (e.g. basal to ciliated prior to birth (Daniely et al., 2004), goblet cell development (Boucherat et al., 2013; Engelhardt et al., 1995; Evans et al., 2004; Hayashi et al., 2004; Pardo-Saganta et al., 2013; Reader et al., 2003; Turner et al., 2011; Tyner et al., 2006). Additionally, the cell types present in different airway locations vary between mice and humans: club cells are present in the trachea and bronchi of mice but only in bronchioles in humans (Plopper et al., 1980a; Plopper et al., 1980b, c), and basal cells are only present in trachea of mice but extend to both large and small airways in humans (Hogan et al., 2014; Rock et al., 2011; Rock et al., 2009), reviewed in (Lynch et al., 2019; Tata and Rajagopal, 2017). Even less well understood is the contribution of rare cells such as ionocytes to small airway epithelial function. A recent study by Okuda et al found no ionocytes (defined as FOXI1-positive, CFTR-expressing cells previously detected in mouse and human airways) in human terminal bronchioles, which has important implications for CFTR function (Okuda et al., 2020).

The porcine lung is a compelling model to study the composition and function of small airway epithelia. Pig and human airways have similar lung anatomy (Judge et al., 2014) and similar distribution of basal, ciliated, secretory (including club) cells along the airway tree when assessed by histopathology (Plopper et al., 1980a; Plopper et al., 1980b, c). Additionally, the pig and human immune system are closer in function and structure than that of mice (reviewed in (Pabst, 2020)). Finally, *CFTR-*deficient pigs develop lung disease mimicking human cystic fibrosis (Meyerholz et al., 2010; Pezzulo et al., 2012; Rogers et al., 2008; Stoltz et al., 2010).

Here, we studied the key differences in large and small airway epithelial composition and gene expression and hypothesized that the gene expression profile of a given cell archetype (Montoro et al., 2020; Schupp et al., 2020) would depend on its location. We used airway microdissection to isolate freshly-obtained small airway epithelial cells (Li et al., 2016) which in newborn pigs correspond to those less than 200 μm diameter. We then performed a single cell resolution census of large and small pig airways, and compared wild-type to *CFTR*^-/-^ pigs to determine the effect of CFTR on epithelial composition and gene expression. We investigated the relative contribution of different cell archetypes in the small airways to ion transporter gene expression, and compared the expression profile of these cells to their corresponding large airway cell archetype. We found that whereas the small and large airways share similar major cell archetypes, their transcriptional state is highly determined by location and shows major differences that correlate with cellular function.

## Results

### Study design

Details on tissue dissection, single cell suspension, library construction, sequencing, and bioinformatics analysis can be found in **Materials and Methods**. Large and small airway tissue cells from lungs of multiple newborn *CFTR^+/+^* and *CFTR^-/-^* pigs were sequenced; single cell RNA-seq analysis was performed summarizing data at the subject level to increase statistical rigor.

### Common cell types are present in both large and small airway epithelia

We sequenced 8,928 large airway cells and 17,773 small airway cells, both shown in **Figure 1**. Our clustering analysis identified 10 major cell types (**Figure 1A**), including epithelial and non-epithelial cells. **Figure 1B** shows the top cluster-enriched genes for the ten cell types. **Supplemental table 1** contains averaged log transformed gene expression for each cluster identified. We quantified the relative abundance of each cell type in epithelial and non-epithelial subgroups (**Figure 1C**) to measure the composition of airway tissues. We initially analyzed epithelial cells and predicted that ciliated cells would be less abundant in the small airways, whereas other common cell types would be more abundant (Boers et al., 1998, 1999; Plopper et al., 1980a; Plopper et al., 1980b, c). Among epithelial cells, the majority were ciliated cells (92.6% in large and 66.2% in small airways, consistent with our hypothesis), with fewer basal (4.9% in large, 21.2% in small airways) and secretory cells (1.8% in large, 11.9% in small airways). There were rare cells (<1% in large and small airways) with high expression of tuft, ionocyte, and alveolar progenitor cell marker genes. The tuft (brush) cells and ionocytes were part of the same main cluster, so for subsequent analysis, we split this group into two subgroups based on whether *FOXI1* and *CFTR* were co-expressed (“Ionocytes”) or not (“Rare”); rare cells that cannot be clearly assigned an ionocyte or tuft (brush) identity were also recently identified in human tissue samples (Deprez et al., 2020; Ruiz Garcia et al., 2019). In addition to epithelial cell types, we also found large numbers of non-epithelial cells (**Figure 1C**), including fibroblasts, endothelial cells, immune cells, and smooth muscle cells in both large and small airway samples. The method utilized for tissue processing large and small airways differed and may bias the large airway samples towards surface cells; reassuringly, the presence of non-epithelial cells in both large and small airway samples shows that the full thickness of the epithelium was reached. Moreover, we sampled enough cells from each major cell type to perform statistically robust comparisons.

**Figure 1:**
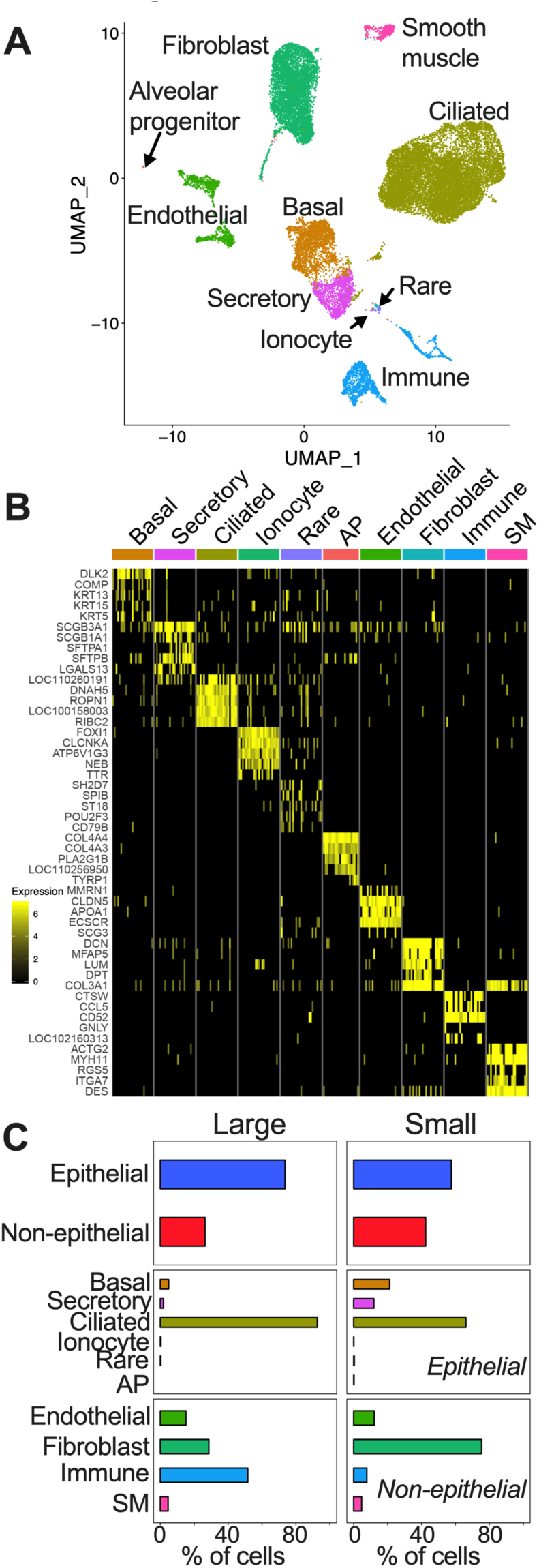
Single cell RNA-seq of large and small airway tissue from *CFTR*^+/+^ and *CFTR*^-/-^ pigs reveals ten cell types. (A) UMAP cluster visualization. (B) Marker genes heatmap (C). Cellular composition in large and small airways. SM = Smooth muscle, AP = Alveolar progenitor. n = 10 large and 7 small airways.

Immunofluorescence confocal microscopy (**Supplemental figure 1**) confirmed that the surfaces of both large and small airways are predominantly lined by acetylated α-tubulin(+) ciliated cells. Both the small and large airway surface included SCGB3A2(+) and MUC5AC(+) secretory cells. KRT5(+) basal cells were also detected in both large and small airways. These data show that small and large airways share the same common epithelial cell types although their relative abundances may vary.

### Ionocytes were not detected in the small airways

Ionocytes are a rare airway epithelial cell type with an expression pattern resembling renal intercalated cells (Montoro et al., 2018; Plasschaert et al., 2018; Scudieri et al., 2020; Vidarsson et al., 2009); they likely participate in airway epithelial ion and fluid transport. Airway ionocytes express the transcription factor *FOXI1*, very high levels of *CFTR*, and genes for V-ATPase subunits. We found a small cluster of ionocyte-like cells in our single cell RNA-seq data based on high average expression of various previously described ionocyte markers. First, we identified cells co-expressing *CFTR* and *FOXI1* in wild-type pig single cell RNA-seq data (**Figure 2A**). We found 2 to 6 ionocytes per 1,000 cells in large airways and none in the small airways. Second, we used protein expression of Barttin (BSND) (Rehman et al., 2020) measured by immunocytochemistry to identify ionocytes in large and small airway tissue. We identified BSND (+) cells in large but not in small airways (**Figure 2B**). Given the absence of CFTR-rich ionocytes in small airways, we expected that *CFTR* would be expressed at lower overall levels in small airways, or *CFTR* expression would be similar in small airways with other epithelial cells compensating for the loss of ionocyte *CFTR*.

**Figure 2:**
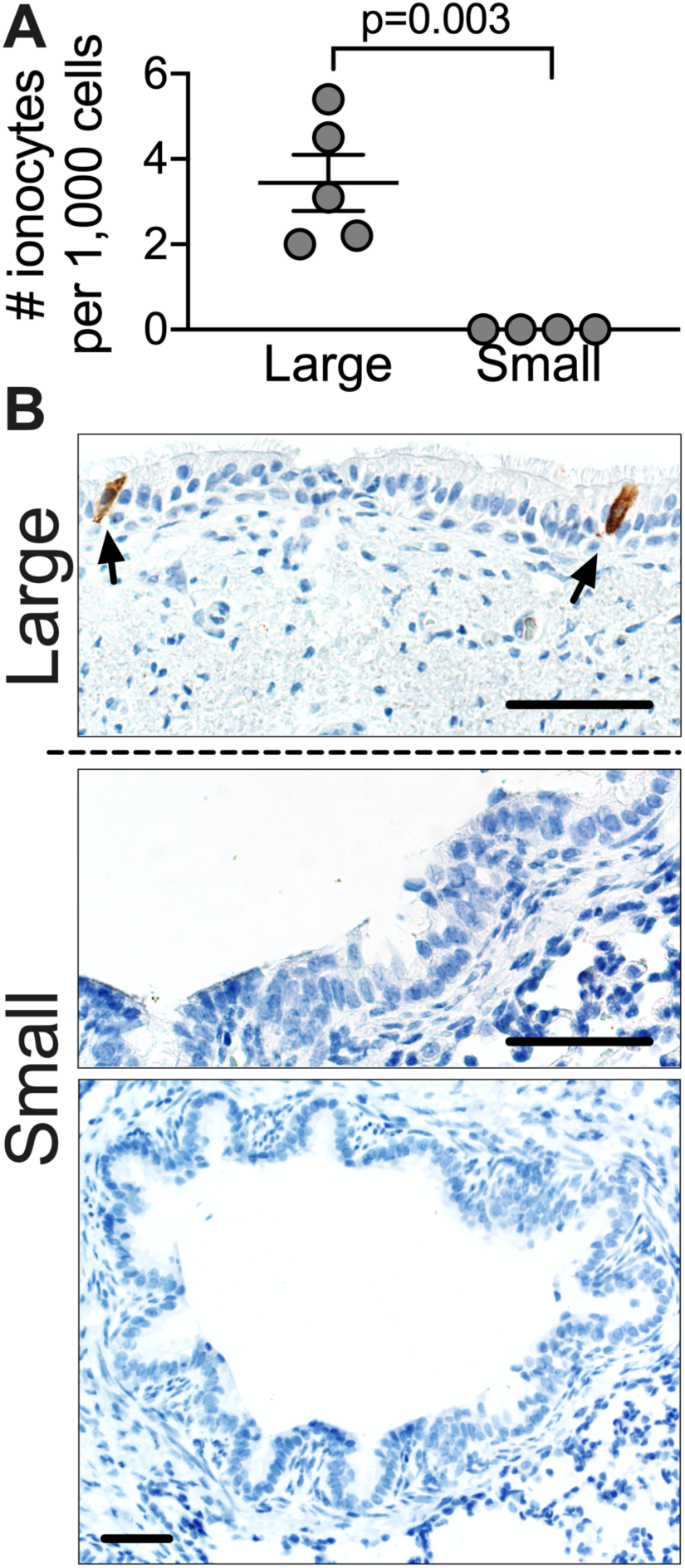
Ionocytes were not detected in small airways of *CFTR*^+/+^ pigs. (A) *CFTR+/FOXI1+* (ionocyte) cell frequency per single cell RNA-seq in large and small airways. (B) BSND immunohistochemistry staining of small and large airway epithelia. Scale bar = 75μm. n = 5 large and 4 small airways.

### Increased expression of *CFTR* transcripts in secretory cells in the small airways

To characterize the pattern of *CFTR* expression in large and small airway epithelia, we determined the fraction with detectable *CFTR* **(Figure 3A)** and the average *CFTR* expression level **(Figure 3B and Supplemental table 1)** for each cell type. Since *CFTR* is expressed at relatively low levels in most cell types yet background transcripts may contaminate single cell RNA-seq data, we show a cell type (endothelial cells) in which *CFTR* transcripts were rarely detected for comparison.

**Figure 3:**
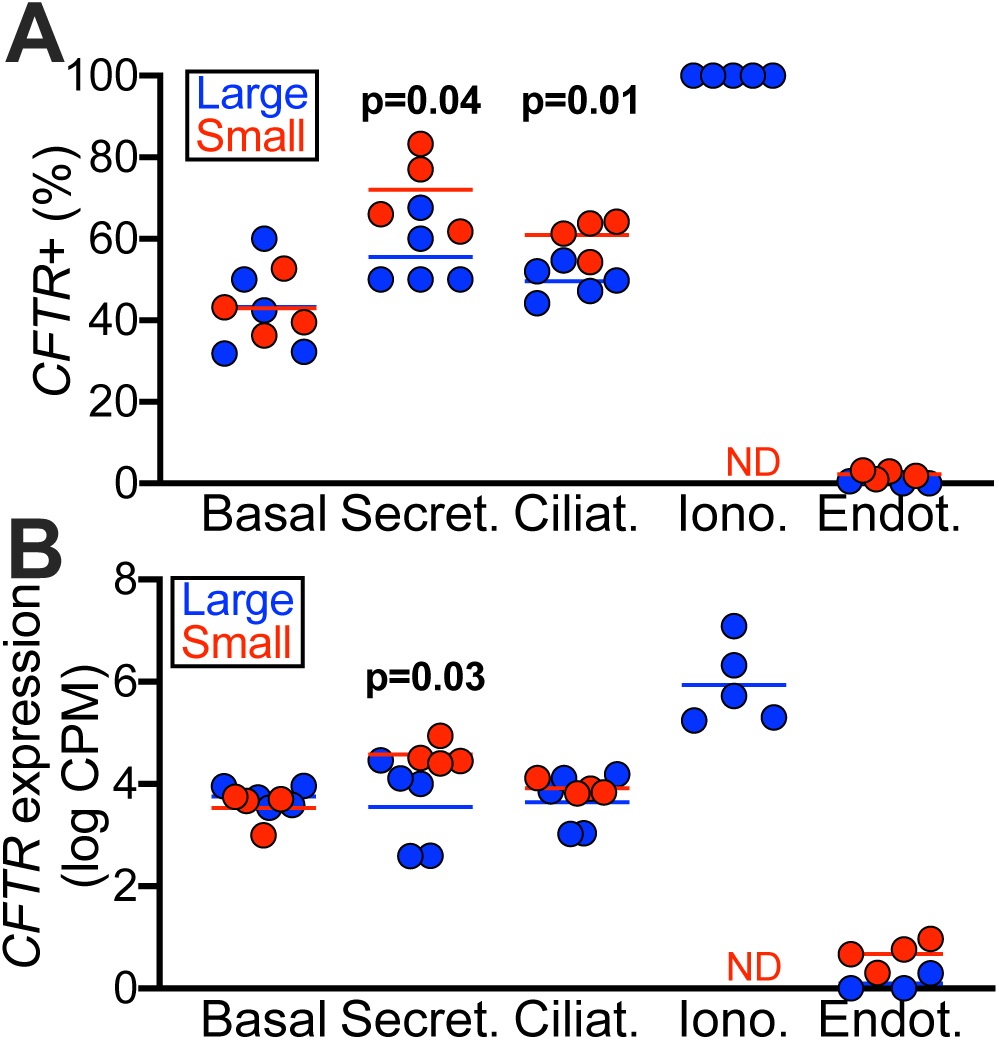
***CFTR* expression in airway surface epithelia of *CFTR*^+/+^ pigs.** (A) percentage of cells with *CFTR* count greater than zero, and (B) log-transformed (counts per million) expression of *CFTR*. ND = not detected. n = 5 large and 4 small airways.

*CFTR is most often detected in ionocytes* **(Figure 3A)**: Detection in ionocytes was 100% by our definition (*CFTR* and *FOXI1* co-expressed). Among the other cell types, *CFTR* was detected most often in secretory cells (50-70%) followed by ciliated, and lastly basal cells. Of note, this fraction of detection is similar to that observed using single cell RNA in situ hybridization and single cell quantitative PCR in human cells (Okuda et al., 2020)

*Secretory and ciliated cells express CFTR more frequently and at higher levels in the small than large airway* **(Figure 3A and 3B)**: We measured increased detection of *CFTR* in secretory and ciliated cells in small airways compared to large airways (secretory: 72±4.9% vs 55±3.5% respectively, ciliated: 61±2.3% vs 49±1.8% respectively). We also detected increased *CFTR* expression level in small airway secretory cells compared to large airways (4.6±0.12 vs 3.6±0.4 Log_2_CPM, respectively).

These data show that *CFTR* expression in the large airways occurs at two levels, with very high expression in a small number of specialized rare cells, and lower expression in the broader population. In contrast, small airway *CFTR* expression was more widely distributed.

### Lack of *CFTR* has a minimal effect on gene expression of common airway epithelial cell types

We and others have previously shown overall minimal gene expression differences in bulk RNA-sequencing between large airway epithelia from people with and without cystic fibrosis and in the wild-type vs. *CFTR*^-/-^ pigs (Bartlett et al., 2016; Pezzulo et al., 2012; Zabner et al., 2005). We predicted that lack of *CFTR* would affect gene expression in a specific cell type in a manner that would not be detectable by bulk RNA-seq. We compared large and small airways from wild-type (*CFTR^+/+^*) and *CFTR^-/-^* pigs. We stratified cells by cell type (basal, secretory, ciliated) and airway type (large and small). For a given cell type, gene counts were aggregated across all cells from individual biological samples to account for variation in gene expression between subjects using the Bioconductor package aggregateBioVar (Huber et al., 2015; R Development Core Team, 2016; Ratcliff, 2020). We used DESeq2 (Love et al., 2014) to perform the differential expression test. The results are shown in **Supplemental figure 2**.

We found less than four differentially expressed genes between *CFTR^+/+^* and *CFTR^-/-^* pigs for any cell type or region. The only consistent difference in gene expression across multiple cell types and airway regions was *CFTR* itself, as we would expect in a comparison of knockout and wild-type animals. These data suggest that, in line with previous observations, lack of CFTR activity minimally modulates expression of other genes in airway epithelial cells in newborn pigs.

### Cell type-specific gene expression varies geographically in airway epithelia

Cells with similar morphology and canonical gene expression biomarkers are generally assigned the same cell identity, but their gene expression profile and physiology may vary because: 1) transcriptional regulation is modulated by the cellular microenvironment, with cells having a location-dependent gene expression profile, or 2) the developmental origin and history of two seemingly identical cells may differ, which would likely affect their expression profile and function (Fu et al., 2017; Morrisey et al., 2013; Rao et al., 2020; Wang et al., 2019; Wang et al., 2015; Xian et al., 2018; Zuo et al., 2015).

We found that for any given cell type, large and small airway cells clustered together in a dimensionality-reduction visualization (**Figure 1A)** and expressed markers that correspond to morphologically defined cell types (**Figure 1B)**. To determine whether expression of genes other than cell-type biomarkers varies geographically, we compared the transcriptional profile of all cell types identified in the large airways to their corresponding cell types in small airways.

#### Small and large airway common epithelial cell types show site-specific gene expression

We expected that a few genes would be differentially expressed in large and small airways based on prior literature and bulk RNA-seq data (Li et al., 2016) including aquaporin 4 (*AQP4*), Surfactant protein D (*SFTPD*) and integrin α 9 (*ITGA9*) expected to be expressed largely in small airways. We were surprised by the large magnitude of differences between corresponding cell types in small and large airways. **Figure 4 (top panels)** shows the results of differential expression analysis. **Supplemental table 2** contains the summarized data for differential expression and marker gene analysis. We discovered 406 differentially expressed genes (Bonferroni adjusted p-val <0.05 and log_2_ fold change <-1 or >1) between small and large airways in basal cells, 746 in secretory cells, and 2,546 genes in ciliated cells. While we found large expression differences in common cell types in the large vs small airways, this was not the case for cell types localized deeper from the epithelial surface such as endothelial and smooth muscle cells (**Supplemental figure 3)**.

**Figure 4:**
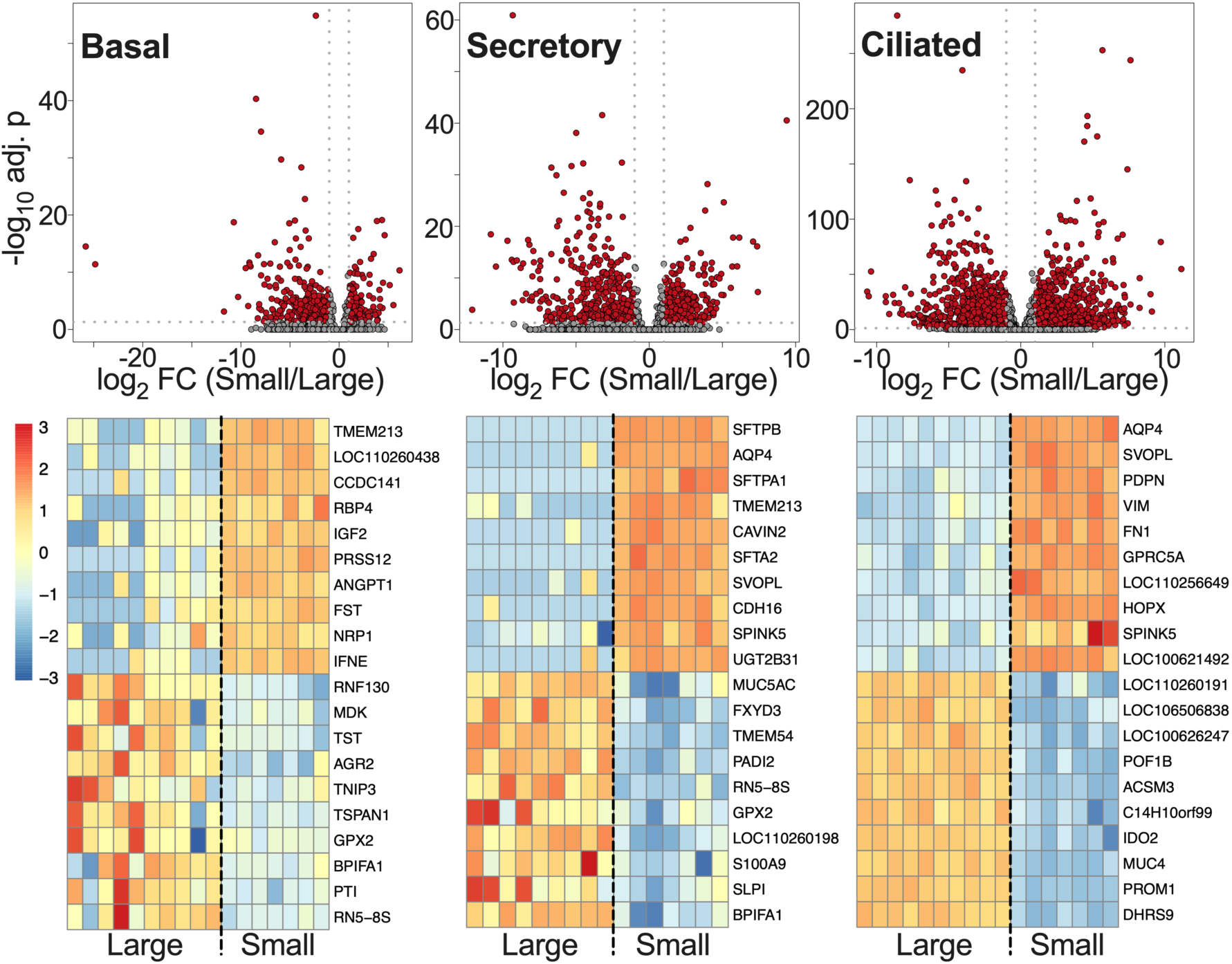
Cell type-specific gene expression varies between large and small airways. Volcano plots and top twenty cell-specific differentially expressed genes in basal, secretory, and ciliated large and small airway epithelia. n = 10 large airway, 7 small airway. *CFTR*^+/+^ and *CFTR* ^-/-^ pig samples were grouped together.

#### Most site-specific gene expression is also cell type-specific

We then investigated whether differential gene expression in large vs. small airways is cell type-specific or shared among the three main epithelial cell types (**Supplemental table 2)**. We used a cutoff of log_2_ fold change 1 for differential gene expression between large and small airways and false discovery rate < 0.1 for genes detected in at least 30% of basal, secretory, or ciliated cells; genes passing these filters were clearly above background levels of detection. Using these criteria, we found 384 differentially expressed genes in large vs. small airways in basal cells, out of which 118 were differentially expressed only in basal cells. A total of 849 genes were differentially expressed in secretory cells, with 440 exclusively in secretory cells. In ciliated cells, 787 genes were differentially expressed, of which 457 were exclusive to ciliated cells. Therefore, roughly half of differentially expressed genes were specific to a cell type. **Figure 4 (lower panels)** shows the top 20 up- and down-regulated genes in small vs. large airways for the three major epithelial cell types.

Only 113 shared genes were differentially expressed among all three main cell types (**Supplemental table 2 - Epithelial intersection**). Genes expressed at higher levels in all large airway epithelia common cell types included well-characterized genes involved in goblet cell metaplasia including anterior gradient 2 (*AGR2*) which is a protein disulfide isomerase involved in the epithelial allergic response and mucin production (Schroeder et al., 2012), and chloride channel accessory 1 (*CLCA1*) which participates in MAPK signaling driving mucus expression and in TMEM16A-mediated chloride transport (Nagashima et al., 2016; Sala-Rabanal et al., 2015).

#### Small airway basal cells express barrier-forming claudins and apical membrane ion transport genes and reach the apical surface

We were surprised by the high levels of expression of genes coding for proteins normally important for transepithelial ion transport in small airway basal cells, as basal cells are not thought to participate in apical-basolateral ion transport. This was particularly striking for genes such as the amiloride-sensitive sodium channel (ENaC) *SCNN1B* and *SCNN1G* which are considered apical surface sodium channels in airway epithelia(Chambers et al., 2007; McDonald et al., 1995; Smith et al., 1994; Smith and Welsh, 1993; Snyder et al., 1994; Zabner et al., 1998), and the barrier-forming claudin 1 (*CLDN1*) **(Supplemental tables 1 and 2) (Flynn et al., 2009; Gan et al., 2013)**; these three genes were among the top 20 genes highly expressed in small vs. large airway basal cells. Based on the expression of genes typically considered important for function of apical surface epithelial cells, we hypothesized that basal cells in the small airways reach both the basement membrane and the apical surface.

We performed a detailed examination of the abundance and localization of basal cells in small and large airway epithelia. Tumor protein 63 (p63) is a well characterized basal cell marker (Daniely et al., 2004; Fernanda de Mello Costa et al., 2020; Hawkins et al., 2021; Rao et al., 2020; Rock et al., 2009; Suprynowicz et al., 2012; Zuo et al., 2015). **Figure 5A** shows that in the distal respiratory bronchioles, which contain mostly a monolayer of cuboidal epithelial cells, there were clear p63^+^ cells that contact the lumen similar to all cells that surround them. Interestingly, some p63^+^ cells also appeared to have cilia, which contrasts with the prior evidence that basal cells first differentiate into secretory cells prior to differentiation into ciliated cells in the large airways.

**Figure 5:**
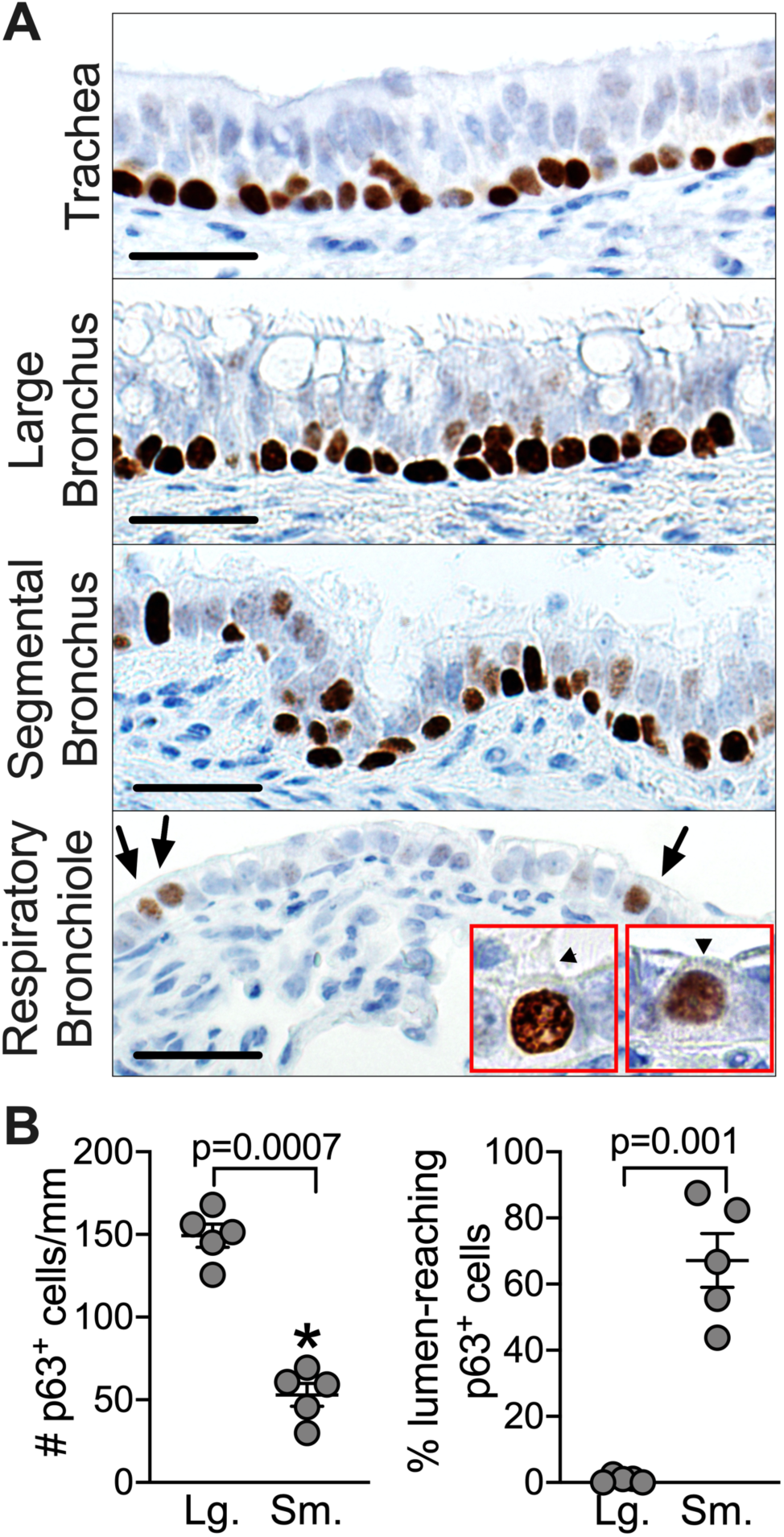
Small airway basal cells are surface cells. (A) immunohistochemistry (40X, scale bar = 25μm) at various airway levels. Arrows: surface p63^+^ cells, red inset (100X magnification) surface p63^+^ cell with and without visible cilia. n = 5 large and 4 small airways. (B) immunohistochemical quantification of basal cells and their position in large and small airways.

Transcripts for *TP63* were clearly detected by single-cell RNA-seq in basal cell clusters. Using immunohistochemistry to detect p63^+^ cells **(Figure 5B)**, we found that while small airway epithelia had less p63^+^ cells per surface unit, most small airway p63^+^ cells contacted both the basement membrane and the airway lumen. In contrast, almost none of large airway basal cells contact the lumen. Moreover, 5.5±1% of small airway basal cells had an expression profile consistent with the mitotic state required for progenitor cells, suggesting small airway basal cells may, like large airway basal cells, function as stem/progenitor cells for other small airway cell types.

#### Small airway secretory and ciliated cells have a distinct mucin gene expression profile

Given their importance for airway physiology in health and disease, we examined expression of mucin genes in large and small airways (**Figure 6A, Supplemental table 1, Supplemental figure 4**) (Okuda et al., 2019). Expression of the secreted mucins MUC5AC and MUC5B was site-specific. In particular, we observed the previously described transition of secretory cells from goblet-like phenotype (e.g., *MUC5AC*-rich) proximally in large airways to club-like phenotype (e.g., *MUC5AC*-low) distally in small airways (Bonser and Erle, 2017; Widdicombe, 2013; Zuo et al., 2018). *MUC5B* expression was also higher in the large airways. Small airway secretory cells expressed higher levels of tethered mucins *MUC1* and *MUC15,* and lower levels of *MUC13* and *MUC20* compared to large airways. Together with differential expression of goblet cell genes *CLCA1* and *AGR2*, these data suggest an upstream site-specific expression program promoting secreted mucins in the large airways.

**Figure 6:**
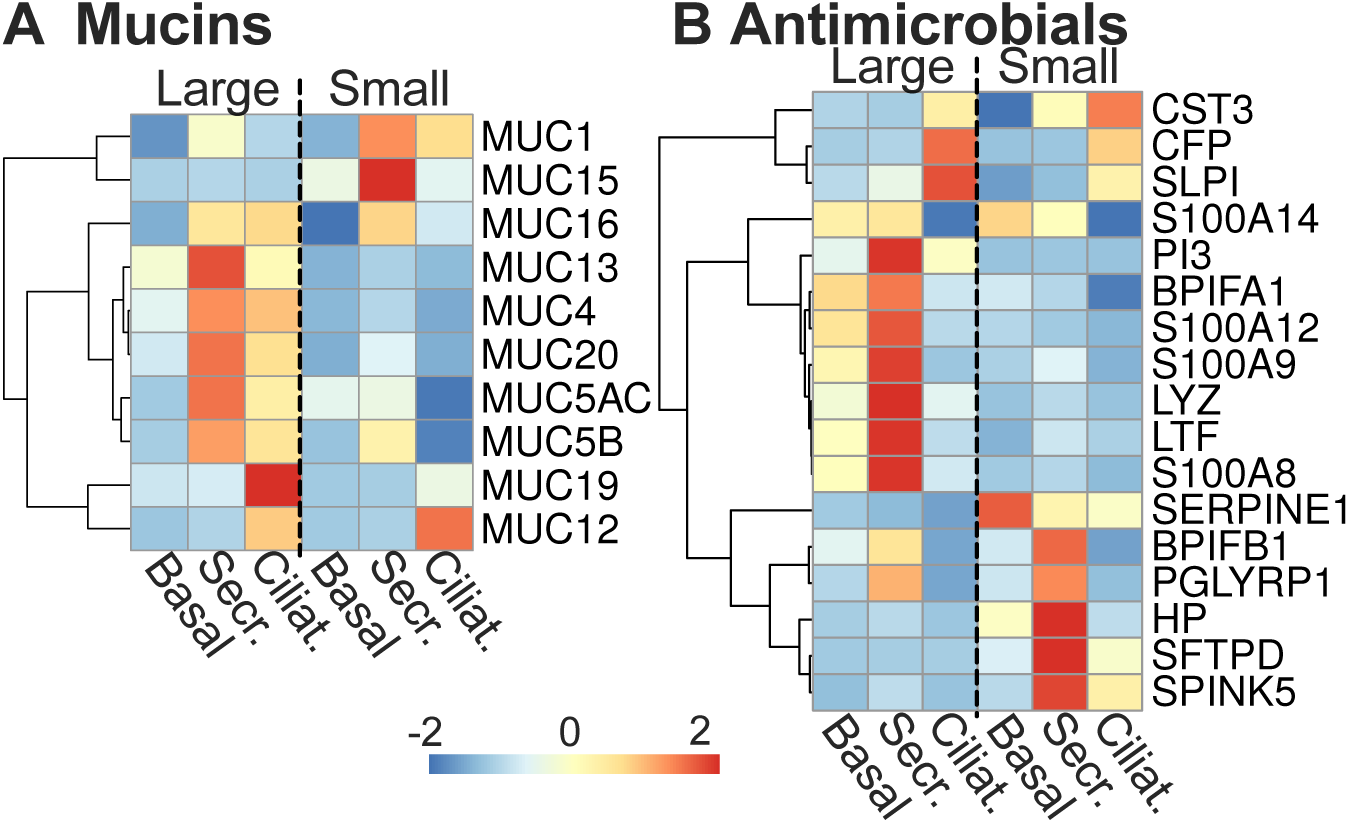
Mucin and antimicrobial genes are differentially expressed between large and small airway surface epithelial cells. (A) Mucin and (B) antimicrobial genes heatmaps: normalized/centered average gene expression in large and small airway for cell types indicated, genes detected at 2 or higher TPM. n = 10 large and 7 small airway.

#### Airway antimicrobial and surfactant gene expression varies in the large and small airways

The airways are constantly exposed to inhaled and/or aspirated microorganisms and depend on various innate and acquired immune mechanisms to maintain near-sterility. Secreted airway antimicrobial molecules can act synergistically to kill microorganisms and are modulated by the local concentration of ions and other factors (Abou Alaiwa et al., 2014; Bals, 2000; Dubin et al., 2004; Grubor et al., 2006; Pezzulo et al., 2012; Simonin et al., 2019). Since large and small airways may be exposed to different amounts and types of bacteria, fungi, and viruses, we expected expression of secreted antimicrobials to be different in these two sites (Sutherland et al., 2010). Overall, secretory cells expressed most antimicrobials (**Figure 6B**). Our data show that compared to small airways, large airway secretory cells express higher levels of lysozyme (*LYZ*), lactoferrin (*LTF*), and calprotectin genes (*S100A8*, *S100A9*, and *S100A12*). In contrast, small airway secretory cells express high levels of the antimicrobial surfactant protein D (*SFTPD*), plasminogen activator-inhibitor 1 (*SERPINE1*), peptidoglycan recognition protein 1 (*PGLYRP1*), haptoglobin (*HP*) and Serine Peptidase Inhibitor Kazal Type 5 (*SPINK5*). Taken together, these data suggest location-specific regulation of secreted innate antimicrobial molecules in large and small airways.

Surfactants are key regulators of epithelial surface tension and are required to maintain alveolar and airway patency (reviewed in (Weaver and Whitsett, 1991; Whitsett et al., 2010)). BPI Fold Containing Family A Member 1 (*BPIFA1*, also known as SPLUNC) has antimicrobial, surfactant, and smooth muscle signaling functions (Ahmad et al., 2016; Garcia-Caballero et al., 2009; Garland et al., 2013; Walton et al., 2016; Wu et al., 2017) and is expressed at high levels in secretory and ciliated cells of the large airway. Expression of *BPIFA1* in small airway secretory cells is decreased almost 50-fold compared to large airways **(Figure 4 and 6B, and Supplemental table 2)**. Instead, and perhaps as an alternative to *BPIFA1*, small airway secretory cells expressed high levels of the surfactants Surfactant Protein B (*SFTPB)* (500-fold higher than large airways) and Surfactant Protein A1 and surfactant-associated 2 (*SFTPA1* and *SFTA2*) (170- and 60-fold higher respectively), these three genes were among the top 6 differentially expressed genes with the criteria used.

### Ion transporter and water channel gene expression patterns correlate with distinct transepithelial ion conductance in small airway epithelia

Our data show that the expression of the anion transporter *CFTR* is markedly different in small airway epithelia compared to large airway epithelia. Small airway epithelia are devoid of *CFTR*-rich ionocytes and have higher expression of *CFTR* in secretory cells, suggesting distinct mechanisms of transepithelial ion transport and water movement in small airways. We therefore investigated the expression patterns of other ion transporters, ion transporter regulators, water channels, and barrier-forming claudin genes in small airway epithelia.

#### Expression and regulation of the amiloride-sensitive sodium channel differs between large and small airway epithelia

The amiloride-responsive transepithelial voltage of airway epithelia decreases in a proximal-distal manner (Ballard et al., 1992; Chen et al., 2010; Shamsuddin and Quinton, 2012, 2014). We expected that expression of genes coding for the amiloride-sensitive sodium channel could be similar (based on our previous in vitro data (Li et al., 2016)) or differ (based on (Okuda et al., 2020; Van Scott et al., 1987)) in large and small airway epithelia. Small airway secretory, ciliated, and basal cells expressed higher levels than large airways of the amiloride-sensitive sodium channel (ENaC) genes *SCNN1B* and *SCNN1G* **(Figure 7A, Supplemental Figure 5 and Supplemental Table 2)**. Interestingly, this correlated with lower levels of *BPIFA1* (*SPLUNC*) which inhibits ENaC expression and function (Garcia-Caballero et al., 2009; Rollins et al., 2010), suggesting that ENaC might be more active in small airways.

**Figure 7:**
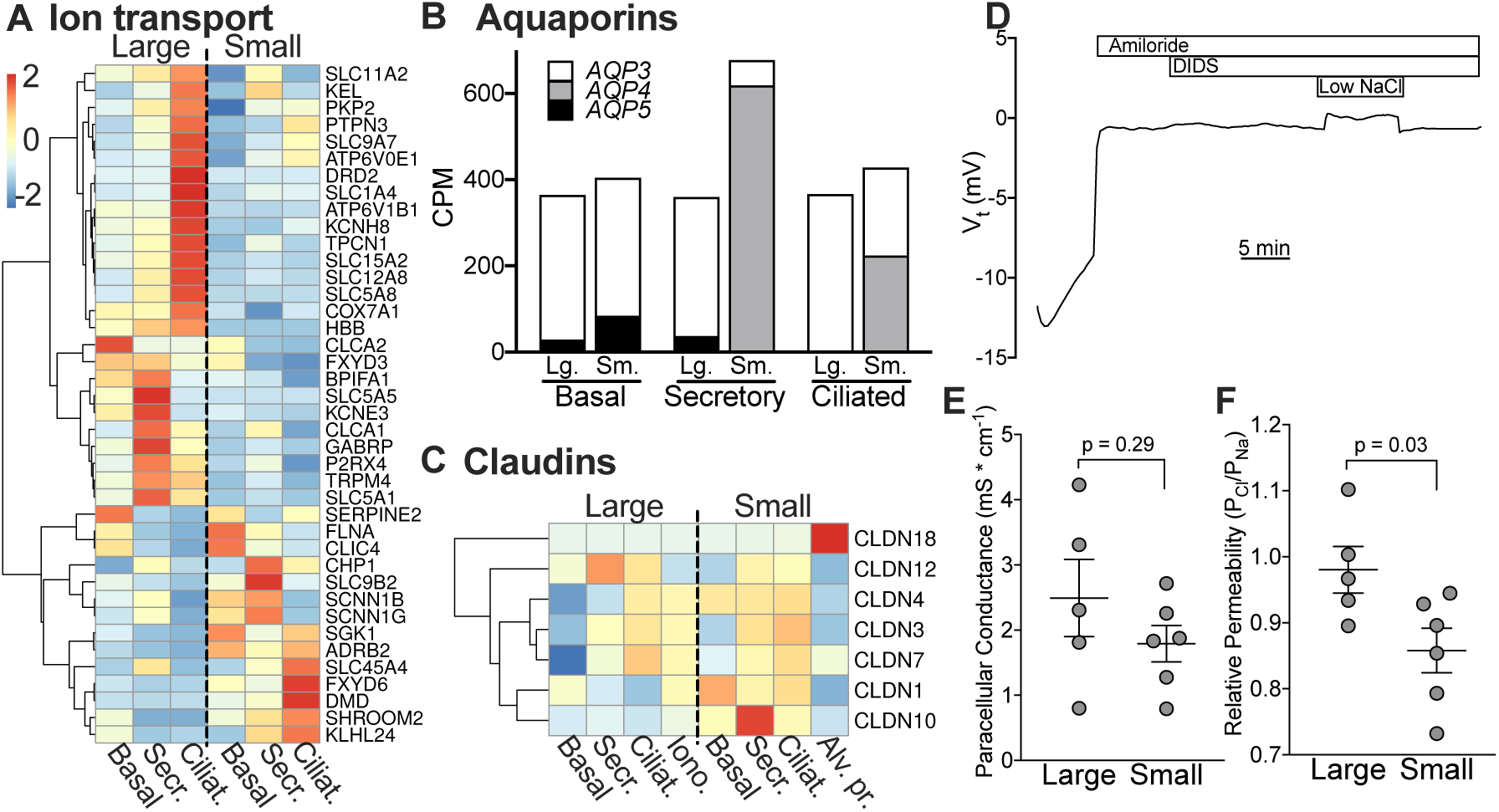
Ion and water transporter gene expression and paracellular ion selectivity differs in small and large cultured airway epithelia. (A) Ion transport and regulator genes heatmap: normalized/centered average gene expression in large and small airway for cell types indicated. (B) Aquaporin 3, 4, and 5 average (CPM) gene expression in common large and small airway epithelia cell types, (C) Claudin genes heatmap: normalized/centered average gene expression in large and small airways for cell types indicated, genes detected at 2 or more TPM. n = 10 large and 7 small airways. (D) Ussing chamber measurements in *CFTR*^-/-^ pig airway epithelia treated with amiloride and DIDS to block transcellular ion transport, (E) shows paracellular conductance, and (F) shows relative paracellular permeability to sodium and chloride. n = 5 large and 6 small airway samples.

#### Expression of genes important for transepithelial water movement differs between large and small airway epithelia

Given the differences in expression of ENaC genes and *CFTR* in small and small airway epithelia, we investigated expression of transcellular water channels and tight junction genes which further modulate transepithelial ion and water transport in polarized epithelia. The expression of aquaporins has been well investigated in large airways and alveoli, but less so in small airway epithelia (Kreda et al., 2001; Verkman, 2007). We determined expression of aquaporins in common large and small airway epithelial cells **(Figure 7B and Supplemental Table 1).** Only aquaporins 3, 4, and 5 (*AQP3, 4,* and *5*, respectively) were consistently detected in at least one cell type. Our data show that *AQP3* is the predominant aquaporin expressed in large airway basal, secretory, and ciliated cells (at 335, 322, and 364 CPM, respectively). *AQP3* is also predominant in small airway basal cells (at 319 CPM) but was expressed at lower levels in small airway secretory and ciliated cells (58 and 204 CPM, respectively). In contrast, small airway secretory and ciliated cells express large amounts (615 and 222 CPM, respectively) of *AQP4*. *AQP4* was one of the top differentially expressed genes in ciliated and secretory cells of the small vs. large airway at near 200-fold higher levels in small airway cells **(Supplemental Table 2)**. Additionally, small airway secretory cells expressed near twice as many total aquaporin transcripts than any large airway epithelial cell type; small airway ciliated cells expressed 20% higher total aquaporin transcripts than large airway ciliated cells **(Supplemental Table 1)**. These data suggest that small airway epithelia are more susceptible to osmotic water movement than large airway epithelia.

Next, we compared the expression of claudins, which regulate tight junction water permeability and ion selectivity in large and small airway epithelia (Flynn et al., 2009; Koval, 2013; Krug et al., 2012; Schlingmann et al., 2015; Soini, 2011; Van Itallie and Anderson, 2004). In the large airways, claudins are primarily expressed by surface ciliated and secretory cells. We found that small airway basal, secretory, and ciliated cells expressed the barrier-forming claudins 1, 4 and 10 (the transcript detected corresponds to isoform 10b) at higher levels than large airway cells (**Figure 7C**). Claudin 12 was expressed at higher levels in large airway secretory and ciliated cells than small airway cells. Of note, claudin 18 was detected only in type II alveolar progenitor cells, consistent with the literature (Kotton, 2018; Sweerus et al., 2017). The differential expression of ion transporters, claudins, and the different cellular architecture of large and small airway epithelia, suggests that regulation of ion transport and barrier function is site-specific.

We have previously shown that cultured small airway epithelia at the air-liquid interface have higher CFTR conductance than and similar ENaC current to large airway epithelia. In contrast, based on our findings in this study, we hypothesized that the paracellular conductance of small airway epithelia at the air-liquid interface in vitro would be different than large airway epithelia. To study paracellular transport, we minimized transcellular ion transport using epithelia from *CFTR*^-/-^ pigs and treated them with the apical ENaC sodium channel blocker amiloride, and 4,4′-diisothiocyanotostilbene-2,2′-disulfonic acid (DIDS) which blocks most non-CFTR Cl^-^ channels in the apical membrane (**Figure 7D**). We then performed Na^+^ and Cl^-^ dilution potential assays and recorded the potential difference that arose by diluting apical NaCl as previously described (Thornell et al., 2020). We found that contrary to our hypothesis, large and small airway epithelia had similar paracellular conductance (**Figure 7E**). However, we found that small airway epithelia had a lower P_Cl_/P_Na_ compared to large airways (**Figure 7F**); these data suggest that the claudin channels in small airway epithelia are more permeable to cations than to anions, whereas large airway epithelia has similar paracellular permeability to cations and anions.

## Discussion

The branching pattern and anatomical configuration of the mammalian airway tree results in a very different environment in the lumen of the large and small airways. In the large airways, laminar cyclic airflow results in respiratory cycle-driven fluctuations in CO_2_ and O_2_ concentration, humidity, and temperature. In contrast, small airways are exposed to a more constant temperature (approaching core body temperature), CO_2_ and O_2_ concentrations, and humidity. Additionally, large airways are constantly exposed to large inhaled particles and oropharyngeal aspirates that may contain large amounts of bacteria, whereas small airway epithelia are more often exposed to small particles and their contents. Therefore, we expected that the cellular composition and gene expression of the large vs. small airways would differ in a manner consistent with functional needs associated with the local microenvironment.

In this study, we performed a single cell RNA-seq census of large and small porcine airway epithelia, using marker-free sampling techniques; we found significant differences between large and small airways at the cellular level and at the transcriptional profile level.

### Large and small airway epithelia vary in cellular composition and cellular configuration

We found that large and small airway epithelia share similar common (basal, secretory/goblet, and ciliated) cell types; in contrast, ionocytes were absent in small airway epithelia. Since ionocytes express very high levels of CFTR in the large airways, they are expected to carry important ion transport functions. We did not detect ionocytes in the small airways using either scRNA-seq or immunocytochemistry.

Since ion transport is critically important in the small airways, how does it differ in the absence of ionocytes? The human airway epithelium contains approximately 303 +/- 20 cells per mm (Korhonen et al., 1969; Pavelka et al., 1976), of which 0.3% are ionocytes in the large airways. Of all cells, approximately 1/3 are basal cells that do not reach the luminal surface; therefore, there are approximately 200 lumen-reaching cells/mm surface with an average diameter 5 μm, of which 0.5% are ionocytes. If we assume uniform distribution of ionocytes on the airway surface, we would expect one ionocyte per luminal surface area occupied by approximately 200 cells, resulting in an estimated distance of 71 μm between ionocytes. We speculate that this configuration is likely inefficient or detrimental in the small airways, as it would result in less than one ionocyte/small airway. Moreover, if ionocyte function involves facilitating secretion of large volumes of fluid, this would likely result in small airway obstruction where ionocytes are present (due to the very small airway diameter). We therefore speculate that lack of ionocytes in the small airways simply reflects the configuration of small diameter tubes in a branching tree, making use of specialized ion transport cells (ionocytes) inefficient or detrimental.

We found important differences in the configuration of cell types in small vs large airway epithelia. Specifically, the ion transporter and barrier-forming claudin gene expression pattern of small airway basal cells led us to discover that at least some small airway basal cells reach the apical surface; we speculate that they participate in transepithelial ion transport. This finding has important practical implications. Gene therapy for lung diseases ideally targets pulmonary stem cells. Airway epithelial basal cells are important progenitor/stem cells and were considered difficult to target via aerosolization given their localization beneath the airway surface. Our data suggest that some small airway basal cells may be directly targeted by gene therapy vectors via aerosolization.

### Cell type-associated gene expression and function is topography-dependent

We found airway common cell types that seemingly share the same gene expression biomarkers and have similar morphology by light microscopy may have very different gene expression profiles when in two different locations. Whereas single cell RNA-seq has previously revealed common “sub-sets” or “cell states” that further sub-classify cell types, the sheer magnitude with hundreds or thousands of differentially expressed genes between corresponding cell archetypes (Montoro et al., 2020; Schupp et al., 2020) in large and small airways was striking.

Our data suggest that location-dependent gene expression in airway epithelial common cell archetypes is associated with the local luminal microenvironment which varies along the airway tree; ciliated, secretory, and basal cells, and to a lesser extent, lung fibroblasts had largely location-dependent gene expression. In contrast smooth muscle and endothelial cell gene expression varied minimally in the large and small airway samples, which may reflect a more uniform microenvironment beneath the surface epithelium along the airway tree.

Overall, we detected higher expression of genes involved in responses to inhaled allergens and pathogens in large airway epithelial cells compared to their small airway counterparts. These included *AGR2*, *CLCA1*, *BPIFA1*, *MUC5AC*, and *MUC5B*. In contrast, we detected higher expression of various ion transporters and water channels (*SCNN1B, SCNN1G, CFTR, AQP4*), barrier-forming claudin genes (*CLDN* genes), and surfactants *(SFTPA1, SFTPB, SFTA2)* in various small airway cell types. Expression of airway antimicrobial peptides/proteins also differed between large and small airway epithelia.

*CFTR* expression followed a strikingly different pattern in the large airways, where it is expressed very highly in ionocytes, followed distantly by secretory and ciliated cells, vs. small airways devoid of ionocytes, where it was expressed most highly in secretory cells, and more highly overall than in large airways. CFTR has previously been shown to have an important role in distal lung water transport (Fang et al., 2002; Li et al., 2012). A recent study by Okuda et al also found that *CFTR* expression in the small airways is higher than in the large airways and most likely to be functional in secretory cells; our sequencing depth facilitated detection of *CFTR* transcripts by sequencing at similar rates as those achieved by in situ RNA hybridization and single cell qPCR in the Okuda et al study (Okuda et al., 2020).

Taken together, our data lead us to speculate that ion and water transport in large and small airway epithelia are mediated by similar cell archetypes, but are finely tuned by differences in gene expression: 1) small airway epithelia express more aquaporins, so changes in regulated ion transport are followed by transcellular fluid secretion or absorption, 2) higher expression of CFTR, whose activation is regulated, and lower paracellular anion conductance gives small airways an advantage for fluid secretion, and 3) higher expression/function of ENaC allows higher Na^+^ absorption, whereas higher relative paracellular cation conductance facilitates paracellular reflux/secretion of Na^+^; this would provide support for higher CFTR-mediated secretion when CFTR activity is high, and higher ENaC-mediated absorption when CFTR activity is low. Overall, the data suggest that small airway epithelia have an advantage over large airway epithelia for both rapid absorption and secretion of ions and water. This is consistent with the notion that small airways need to be “wet enough to be pliable” yet “dry enough to remain patent” as described by Shamsuddin and Quinton (Shamsuddin and Quinton, 2012).

### CFTR does not regulate expression of other genes in common epithelial cells of porcine lungs in vivo

*CFTR* expression varies alongside that of many other genes in different cell types, and gene expression can vary greatly in tissue of diseased *CFTR^-/-^* animal models or in people with cystic fibrosis in the presence of infection and inflammation (Schupp et al., 2020). But does CFTR itself regulate expression of other genes via protein-protein interactions or via ion transport-mediated transcriptional regulation? Our previous data suggested that lack of CFTR did not affect gene expression in animal models in the absence of infection or inflammation (Bartlett et al., 2016; Pezzulo et al., 2012; Zabner et al., 2005). Primary human airway epithelia from people with CF cultured at the air-liquid interface have differential expression of some genes, which may reflect epigenetic modifications induced by inflammation. Our data strikingly show that even when analyzing gene expression at the single cell level, knocking out *CFTR* does not appear to consistently modify gene expression except for *CFTR* itself; this was detected in all common epithelial cell types. Therefore, we propose that whereas *CFTR* is heavily co-regulated with other genes, it does not itself modulate expression of other genes.

### Strengths and limitations of our study

Several factors strengthen our results and interpretations. 1) By using an animal model shown to closely represent human anatomy and function, we were able to obtain fresh tissue from anatomically correct locations. 2) Young *CFTR^-/-^* pigs are not yet largely affected by the consequences of chronic inflammation and infection that affect human CF lungs. Human tissue obtained from explanted lungs has a long explant to library prep time, and obtaining large cell numbers correctly from airways <2mm via bronchoscopic brushing is technically challenging. 3) Direct dissection of small airway tissues avoids over-representation of alveolar tissue, and tracheal surface scraping avoids collection of sub-mucosal gland mucous, serous, and ciliated cells that may have a distinct function and transcriptome but may be difficult to differentiate from corresponding surface epithelial cells. 4) Importantly, by combining single cell RNA-seq with stringent statistical principles implemented in a bioinformatics package developed by our group (Ratcliff, 2020), we avoid large numbers of false positive findings that may occur in single cell RNA-seq analysis when inter-individual variability is not accounted for.

Our study is limited by potential biases in proportional sampling of various cell types, which is a common issue with single cell RNA-seq library prep; due to differential isolation, viability, or lysis-sensitivity of specific cell types, the proportions of cell types obtained may not directly represent the proportions of cell types in the tissue of interest, for example, ciliated cells are over-represented in our large airway samples likely due to the sampling method used, although enough cells of each type for analysis were obtained, including fibroblasts and endothelial cells. Fortunately, by using single cell RNA-seq we are able to appropriately compare gene expression profiles; an approach using bulk RNA-seq may be unable to correct for differential cellular composition between conditions, even when using cell-type inference/deconvolution methods.

A cell with similar morphology and marker gene expression may vary in its transcriptome and function due to its environment, or due to its developmental history (Fu et al., 2017; Morrisey et al., 2013; Rao et al., 2020; Wang et al., 2019; Wang et al., 2015; Xian et al., 2018; Zuo et al., 2015). Whereas our experiments were not designed to investigate the relative contribution of the extracellular environment vs. cell-intrinsic gene expression programs in the small vs. large airway transcriptome, our data show how the function and gene expression profile of a cell archetype varies depending on its cellular microenvironment. Our study highlights that important proximal-distal patterns of cellular composition and gene expression observed in gut and kidney analysis (Brunskill et al., 2008; Chabardes-Garonne et al., 2003; LaPointe et al., 2008; Lindgren et al., 2017; Richmond and Breault, 2010; Wang et al., 2020) also apply to the airways with important implications for lung disease pathophysiology and for therapy of lung disease.

## Supporting information

Main supplemental figures

## Acknowledgments

We would like to thank Linda Powers, Mallory Stroik, Nicholas Gansemer, Christian Brommel, Keyan Zarei, Jason Ratcliff, Michael Chimenti, and The University of Iowa Institute for Human Genetics Genomics Facility for excellent technical support. This work was funded in part by: NHLBI K01HL140261 (A.A.P), Cystic Fibrosis Foundation PEZZUL20A1-KB (A.A.P), Gilead Sciences Research Scholars Program in Cystic Fibrosis (I.M.T.), Cystic Fibrosis Foundation University of Iowa RDP (A.A.P., D.A.S., M.J.W., J.Z.), NHLBI P01HL091842 (A.A.P., D.A.S., M.J.W., J.Z.), NHLBI P01HL051670 (D.K.M., D.A.S., M.J.W., J.Z.), NIDDK P30DK054759 (J.Z.), NHLBI T32HL007638 (G.R.I., M.J.W., J.Z.) and T32GM007337 (G.R.I), and the University of Iowa Physician Scientist Training Program (A.A.P).

## Author contributions

Conceptualization: A.A.P, A.L.T, and J.Z.; Methodology: A.L.T. and W.J.; Formal Analysis: A.A.P, A.L.T, X.L., W.Y., D.K.M., and I.M.T.; Investigation: A.A.P., A.L.T, R.V., X.L., W.Y., S.E.M., D.K.M., G.S.R-I., I.M.T.; Resources: A.A.P., A.L.T., X.L., D.K.M., G.S.R-I., D.A.S., M.J.W.; Data Curation: A.A.P., A.L.T., W.Y.; Writing – Original Draft: A.A.P., A.L.T., J.Z.; Writing – Review & Editing: All authors; Visualization: A.A.P., A.L.T.; Supervision: A.A.P., M.J.W., J.Z.; Funding Acquisition: A.A.P., D.A.S., M.J.W., J.Z.

## Experimental Procedures

### Animals

All protocols for obtaining and use of pig tissue were approved by The University of Iowa Animal Care and Use Committee. The development of the *CFTR*^-/-^ pig model has been previously reported (Rogers et al., 2008). Animals were purchased from Exemplar Genetics (Sioux City, IA). All protocols were approved by the University of Iowa Institutional Animal Care and Use Committee (IACUC). Newborn (within the first 24 hours of life) pigs were sedated with ketamine/xylazine (Akorn) and euthanized with phenobarbital sodium/phenytonin sodium (Euthasol; Virbac). Tissues were sampled from individual animals: five (4 male, 1 female) large airway *CFTR*^+/+^, five (2 male, 3 female) large airway *CFTR*^-/-^, four (3 male, 1 female) small airway *CFTR*^+/+^, and three (2 male, 1 female) small airway *CFTR*^-/-^.

### Tissue preparation and single cell suspension

Newborn piglet lungs were excised immediately following successful euthanasia by sedation and then an overdose of Euthasol. Excised tissue was kept moist using a small volume of DPBS (Gibco #14287080). For large airways, tracheal surfaces were scraped to isolate surface epithelia from submucosal glands and placed in cold 1 x DPBS. Small airway tissue was processed as described in (Li et al., 2016; Thornell et al., 2018); briefly, the airway tree was isolated by carefully combing off the parenchymal tissue, followed by blunt micro-dissection to remove the airway vasculature and any remaining alveolar tissue. Small airways (diameter ≤ 200 µm) were then cut away from the airway tree.

Tissues were centrifuged and resuspended in tissue dissociation medium (pronase 1.4 mg/ml; DNase, 50 U/ml). Tissues were dissociated in 37oC, 5% CO2 in ultra-low attachment cell culture flask (# CLS3815-24EA, Millipore Sigma) for 2 hours and gently shaken every 10 minutes. This procedure was terminated by adding 1 volume of cold 1 x DPBS with 10% fetal bovine serum. Dissociated tissues were filtered with 40-µm tissue strainers to collect cells. Cells were washed with ammonium chloride (0.8%, pH 7.4) for 5 min to eliminate red blood cells and spun down (500 g, 5 minutes), washed with 1 x DPBS twice, and resuspended in 1 x DPBS with 0.4 mg/ml bovine serum albumin (# 74719, New England Biolabs) at approximately 1000 cells/µl. Cell viability was measured by trypan blue staining. Samples with 90% or higher viability were retained for further processing.

### 10X Genomics Single Cell 3’ Library prep

Single cells were isolated and tagged with cell-specific barcodes and unique molecular identifiers (UMIs) using the 10x Genomics Chromium Controller provided by the Iowa Institute of Human Genetics at the University of Iowa with a target of 3,000 cells per single cell suspension. cDNA libraries were prepared using the Chromium Single Cell 3’ Reagent Kits v3 system, and all libraries submitted for sequencing passed 10x Genomics quality control specification. Libraries were sequenced on an Illumina HiSeq 4000 system using 150 bp paired-end reads.

### Raw sequence processing

The cDNA sequences were aligned to pig genome reference Sscrofa 11.1 (Warr et al., 2020). CellRanger 3.0.1 (10x Genomics) software was used to parse cell barcodes and UMIs, align sequences, and generate gene-by-cell UMI count matrices. CellRanger was run twice, using either Ensembl (release 96) (Yates et al., 2020) or RefSeq (O’Leary et al., 2016) gene annotations.

### Porcine reference transcriptome optimization

Whereas the Sscrofa 11.1 genome assembly significantly improved annotation of the porcine transcriptome and distal lung tissue DNA/RNA samples were included for sequencing, airway tissue is underrepresented (Warr et al., 2020). Tissue-specific underrepresentation in a reference annotation may lead to inaccurate RNA-seq sequence alignment; following alignment, some reads appeared to be shifted downstream of 3’ untranslated regions of reference genes under one or both annotations, resulting in loss of reads for quantification.

In order to improve the annotation, we manually checked alignment for more than 6,000 genes expressed in human, pig, or mouse airways. We focused on exons exclusively since CellRanger only requires the annotation of exons as reference. The Sscrofa11.1 RefSeq annotation appeared to provide overall higher agreement with aligned reads than Ensembl. We polished the RefSeq gene annotations using the following modifications: 1) 3’ UTRs were elongated; 2) 3’ UTRs shared by two or more genes were removed; 3) genes from alternative pig genome databases (e.g., mitochondrial genes from Ensembl) or human and mouse homologs were added. We required putative modifications to exist in at least one annotation database among UCSC, RefSeq, Ensembl, and Genbank. Approximately 400 genes were modified. The edited pig genome reference was used to generate gene-by-cell UMI count matrices as discussed above.

### Bioinformatics and statistical analysis

UMI count matrices were inspected for quality before subsequent bioinformatics processing. One large airway and one small airway sample (in addition to samples listed above) were discarded due to uniformly low numbers of UMIs per cell (median UMIs: 2,875 and 2,753, respectively). Among the remaining high-quality samples, we retained only cells with 3,000-50,000 UMIs; 1,000-8,000 detected genes; and less than twenty percent of UMIs expressed from mitochondria. In order to avoid inclusion of submucosal and ductal cells that may have been sampled from the trachea, we excluded cells with non-zero counts of DMBT1, a serous cell marker gene.

We identified cell types according to a standard pipeline for single cell RNA-seq via the R package Seurat version 3.1.1 (Butler et al., 2018; Stuart et al., 2019). First, gene counts for each cell were normalized by total UMIs and multiplied by one million, producing gene expression measurements as UMI counts per million. Next, log-transformed gene expression values were centered and scaled by regressing out the percentage of mitochondrial genes and scores for cell cycle stage. The first 40 principal components of the centered and scaled gene expression measurements were used to obtain cell clusters and uniform manifold approximation and projection (UMAP) coordinates. Cell clusters were associated with cell types by finding the most highly upregulated genes in each cluster and cross-referencing with a list of known marker genes.

Differential gene expression (DGE) analyses were performed with pigs as the units of analysis. We created the Bioconductor (Huber et al., 2015) package aggregateBioVar (Ratcliff, 2020) to facilitate analysis at the subject level. First, the gene-by-cell UMI count matrix containing all cells was divided into multiple matrices, with one gene-by-cell UMI count matrix per cell type. Then for each matrix, UMI counts were summed for each pig resulting in one gene-by-pig count matrix for each cell type. Comparisons of cell proportions and individual gene expression levels were performed using a t-test. Genome-wide DGE analyses comparing small and large airways and *CFTR*^+/+^ and *CFTR*^-/-^ pigs were performed using the R package DESeq2 version 1.22.2 (Love et al., 2014). We identified genes with FDR < 0.05 and at least a two-fold difference in expression as differentially expressed. All statistical analyses were performed using R (R Development Core Team, 2016).

### Resource availability

Further information and requests for resources should be directed to and will be fulfilled by the Lead Contact, Joseph Zabner

### Data and code availability statement

All data are available in GEO (Accession GSE150211. The aggregateBioVar analysis package is available in Bioconductor (Ratcliff, 2020).

### Materials availability

This study did not generate new unique reagents.

### Immunofluorescence

For cell type validation in large and small airways of native tissue, newborn pig lung tissues were fixed with 4% paraformaldehyde, embedding in O.C.T. compound (Tissue Tek by Sakura Finetek, Torrance, CA, USA), cryosectioned into 7 μm sections, and permeabilized in 0.2% Triton X-100. After blocking in super blocking buffer supplemented with 2% BSA, the slides were incubated with the following primary antibodies: 1) mouse anti-acetylated α-tubulin, 1:1,000 (T7451, Sigma, St. Louis, MO); 2) mouse anti-Muc5AC, 1:50 (MA5-12178, Thermo Fisher Scientific, Waltham, MA); 3) rabbit anti-SCGB3A2, 1;100 (ab181853, Abcam Inc., Cambridge, MA); 4) rabbit anti-keratin-5, 1:500 (Cat#: 905501, Biolegend, San Diego, CA); and 5) phalloidin conjugated with Alexa 568, 1:50 (A12380, Thermo Fisher Scientific Waltham, MA). Appropriate secondary antibody (goat anti-mouse-Alexa-Fluor-488, goat anti-rabbit-Alexa-Fluor-488 or 568, 1:500; Invitrogen, Carlsbad, CA) was used. Nuclei were counterstained with DAPI. Images were visualized with an Olympus Fluoview FV1000 confocal microscope with a UPLSAPO ×60 oil lens.

### Immunohistochemistry and scoring for p63

NonCF respiratory tissues (trachea and lung, n=5 pigs) were identified from newborn nonCF pig (<48 hrs of age) and sectioned (∼4 µm) from archival paraffin-embedded blocks. Briefly, tissues were rehydrated through a series of xylene and alcohol baths. Tissues were prepared using antigen retrieval (citrate buffer pH 6.0, 5 min at 125°C, Decloaking Chamber^TM^, Biocare Medical, Pacheco, CA, USA), quenching of endogenous peroxidase (3% hydrogen peroxide, 8 minutes) and blocking of nonspecific stain (Background Buster x 30 minutes, Innovex Biosciences, Richmond, CA, USA). A mouse monoclonal anti-p63 antibody (1:500, MS-1081-P, Neomarkers, Fremont, CA, USA) and followed by a secondary kit (Mouse Envision, DAKO-Agilent, Santa Clara CA, USA), stained with chromogen (3,3′-Diaminobenzidine [DAB], DAB Plus and DAB Enhancer, DAKO-Agilent) and counterstained (Harris hematoxylin x 1 minute). Tissues immunostained for p63 were evaluated and scored by a boarded veterinary pathologist following principles for reproducible tissue scoring (Meyerholz and Beck, 2018). Trachea and small airways (<200 µm lumen diameter) from each animal were evaluated for p63 staining. Parameters evaluated included: basement membrane length of airway wall studied, number of p63+ and p63-epithelial cells and the number of p63+ cells that were in proximate contact with the airway lumen. Summary data for large and small airways were normalized per airway length and evaluated using a pair-test with significance placed at P<0.05.

### Culture of porcine large and small airway epithelia at the air-liquid interface

Isolated primary epithelial cells were seeded onto collagen-coated, semi-permeable membranes (Corning #3413). Primary epithelia cultures were maintained at an air-liquid interface at 37°C in a 5% CO2 atmosphere, as previously described (Zabner et al., 1996). Primary small airway cultures were maintained in Small Airway Growth Media (Lonza, Basel, Switzerland) supplemented with 10ng/mL keratinocyte growth factor (KGF), as described previously (Li et al., 2016). Primary large airway cultures were obtained from The University of Iowa Culture Core and were maintained in USG media. ∼2 weeks prior to experiments, small airway epithelia were switched to USG media and handled in parallel to large airway cultures.

### Electrophysiology studies

Epithelial cultures were mounted in Ussing chambers and bilaterally bathed in a 150mM NaCl solution with 1.2 mM CaCl2, 1.2 mM MgCl2, 5 mM HEPES, and 5 mM glucose; pH 7.40; osmolarity 310 ± 5 mOsm. Transepithelial voltage was monitored under open-circuit conditions through 3M KCl agar bridges connected to an amplifier (VCC-MC8, Physiologic Instruments). To obtain conductance, a periodic ± 5 µA pulse was applied across the epithelia. Transcellular ion transport was inhibited by using CF pig epithelia to eliminate CFTR-mediated Cl–transport, 100 µM apical amiloride to eliminate Na+ transport, and 100 µM apical DIDS to eliminate any remaining Cl–transport. Dilution potentials were generated by diluting the apical NaCl solution from 150 mM NaCl to 75 mM NaCl while maintaining osmolarity with mannitol. Relative permeabilities were calculated using ion activities and the Goldman-Hodgkin-Katz equation as previously described (Thornell et al., 2020).

**Supplemental Table 1:**

Log CPM per cell type average across all samples.

**Supplemental Table 2:**

Differential expression and marker gene analysis for large vs. small airway samples.

