## Supplementary material for "Topography-dependent gene expression and function of common cell archetypes in large and small porcine airways": Main supplemental figures

### Primary supplement

Pezzulo et al. 2021

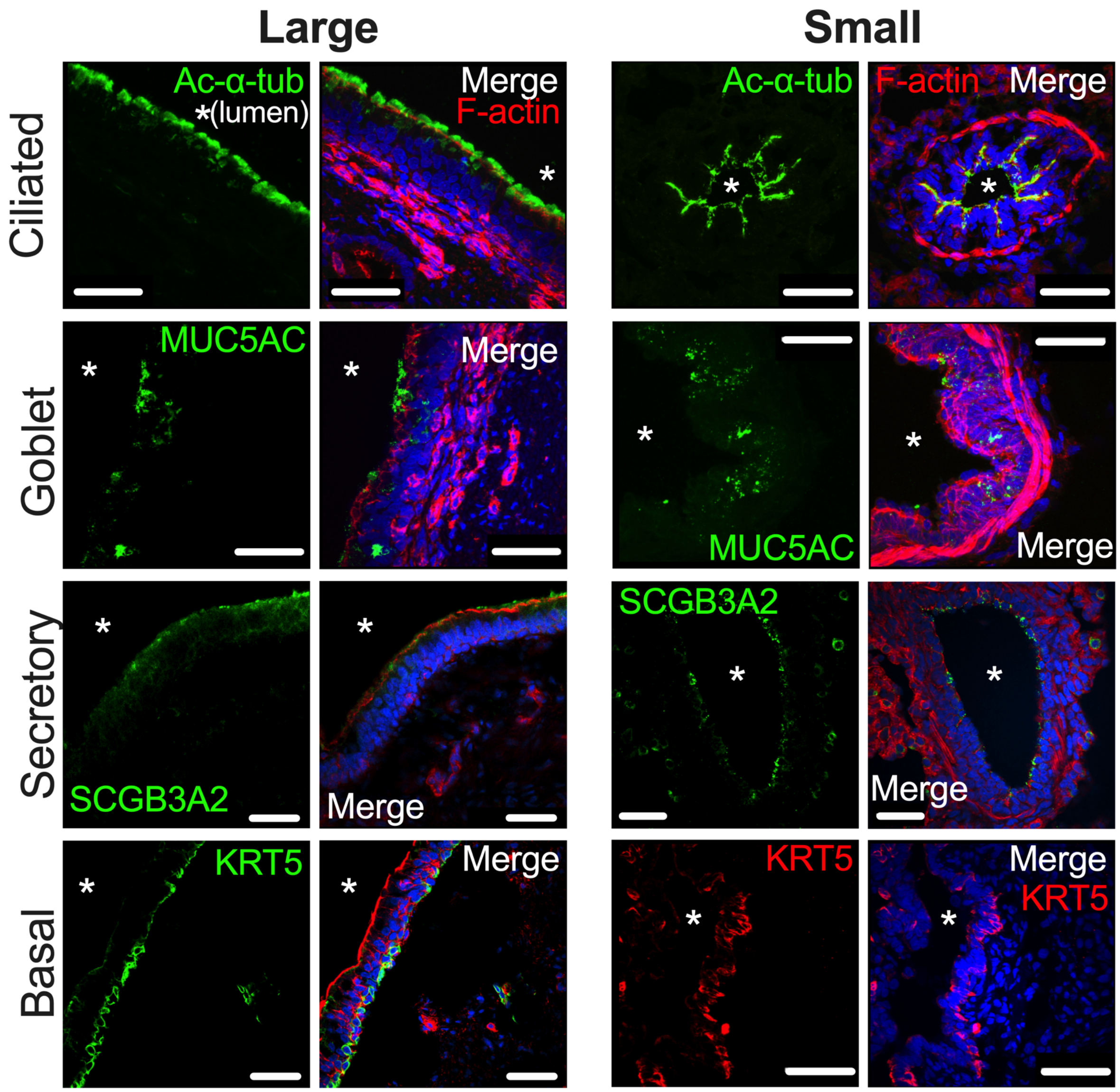


**Supplemental figure 1: Large and small porcine airway epithelia share same common cell types.** Immunofluorescence images showing ciliated (α-acetyl tubulin), secretory (Muc5AC, SCGB3A2), and basal (KRT5) cell markers in large and small porcine airways. Merge images show F actin in red (except for small airway KRT5) and DAPI in blue. Scale bar = 25µm.

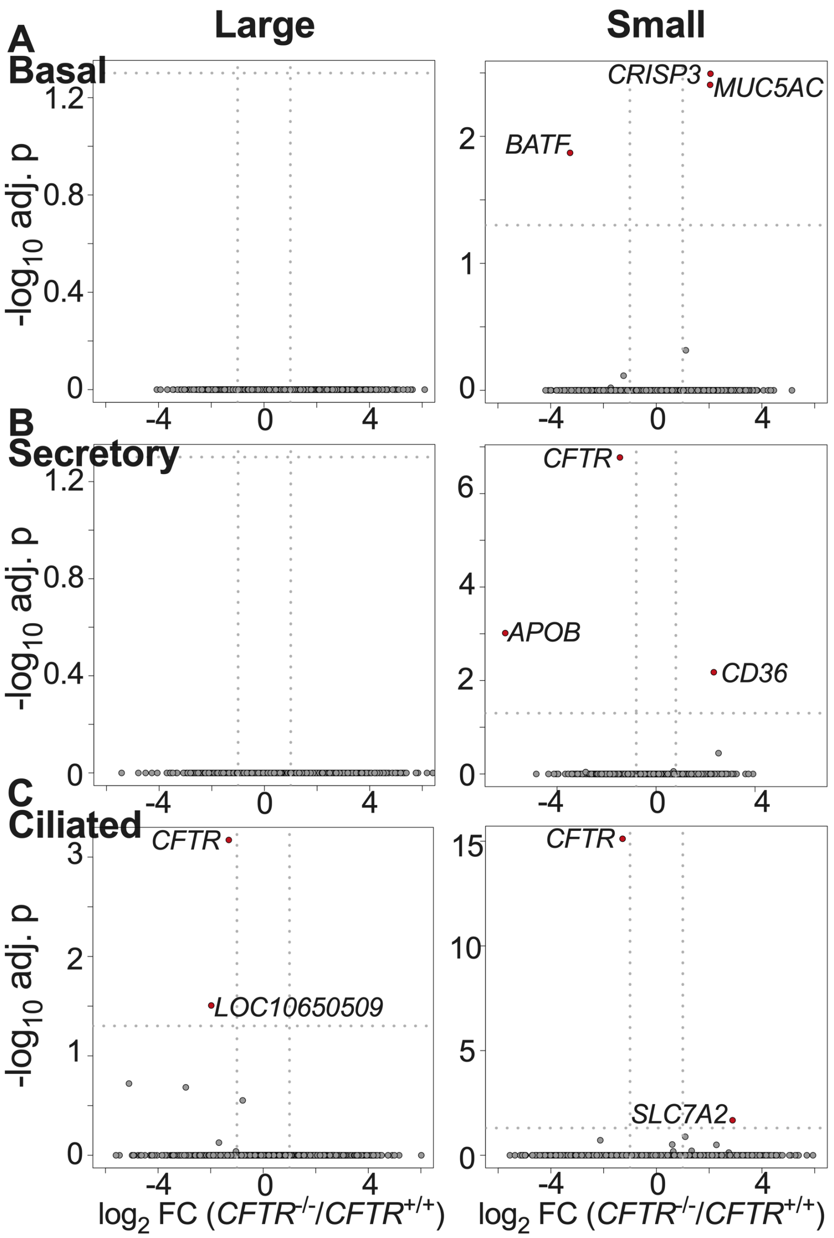


**Supplemental figure 2: Expression profiles of airway surface epithelia from *CFTR*$^{+/+}$ and *CFTR*$^{-/-}$ pigs are similar.** Volcano plots of *CFTR*$^{+/+}$ vs. *CFTR*$^{-/-}$ (A: basal, B: secretory, and C: ciliated) cells in large and small airways. n = *CFTR*$^{+/+}$ pigs (5 large and 4 small airway) and *CFTR*$^{-/-}$ pigs (4 large and 3 small airway)

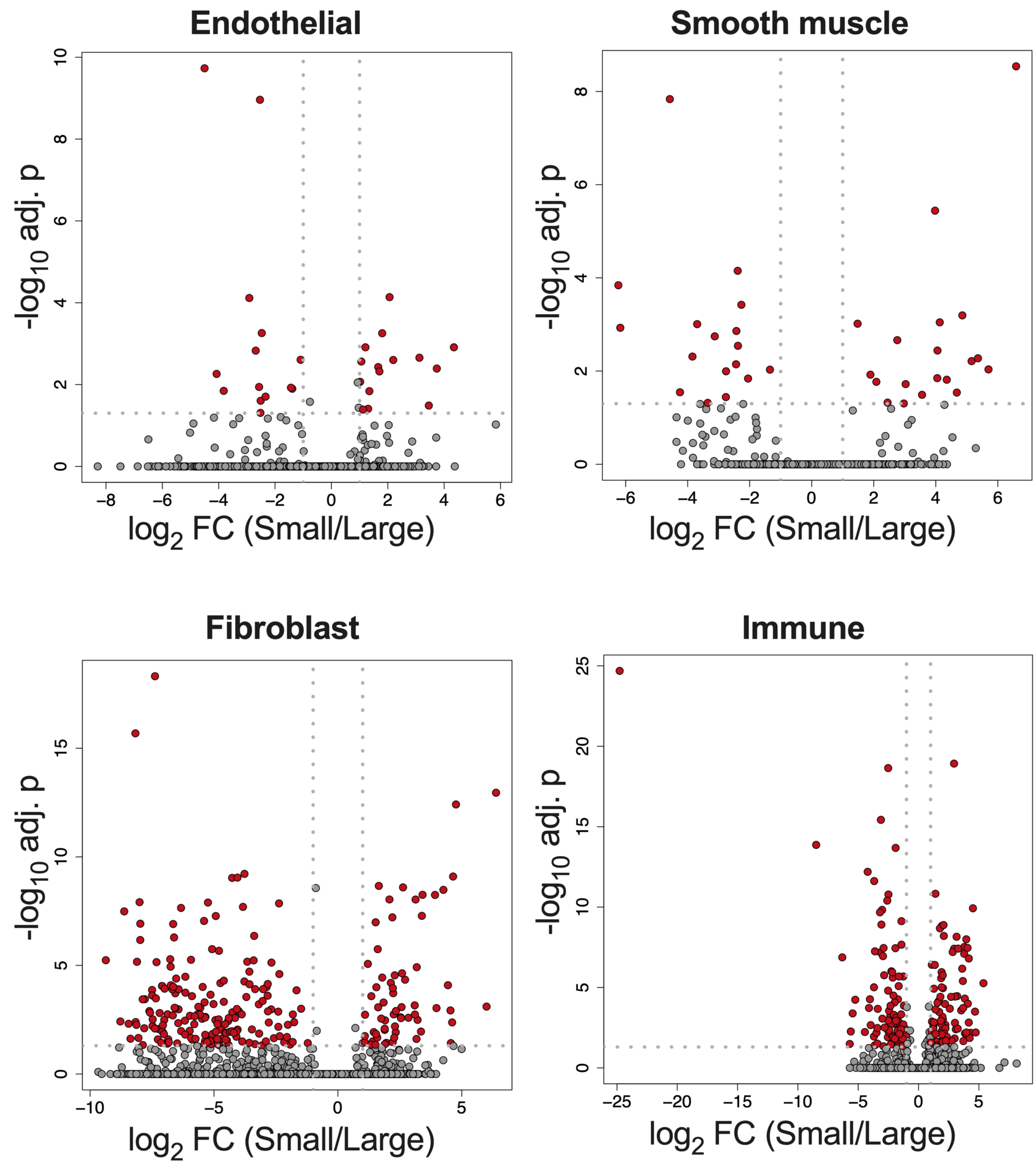


**Supplemental figure 3:** Differential gene expression in non-epithelial cells: Cells that more directly interact with epithelial cells (fibroblasts and immune cells) had more site-specific differentially expressed genes compared to endothelial and smooth muscle cells. n = 10 large airway, 7 small airway. *CFTR*$^{+/+}$ and *CFTR*$^{-/-}$ pig samples were grouped together as they had comparable gene expression.

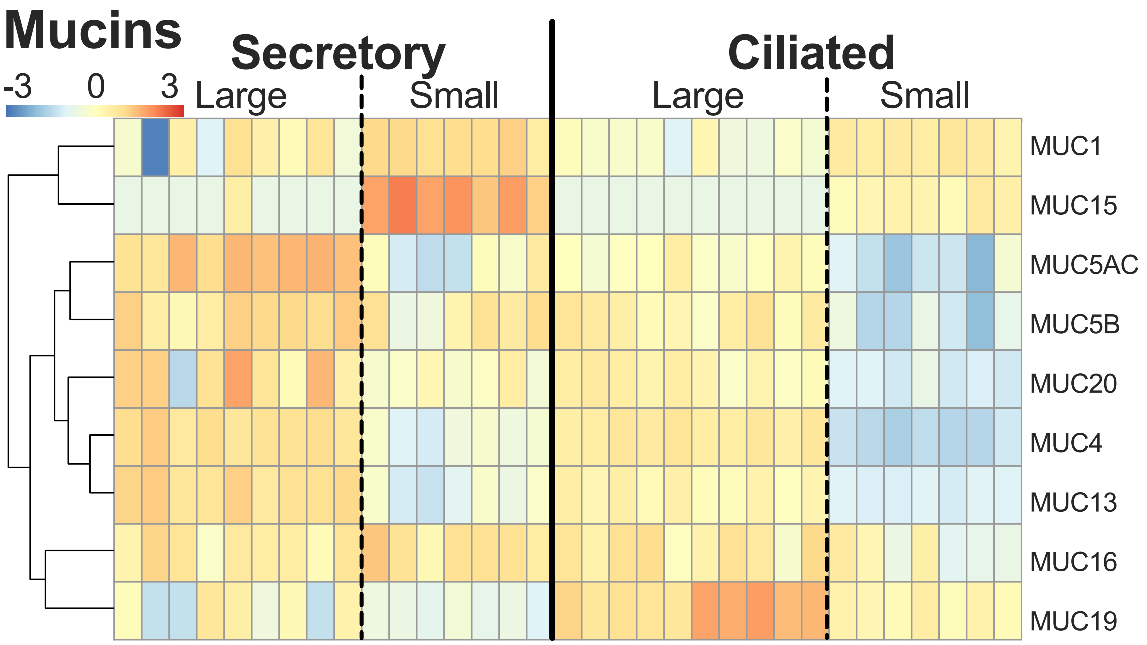


**Supplemental figure 4: Mucin genes expression in small and large airway secretory and ciliated cells at the subject level:** Each column is the normalized/centered average expression per cell type (basal, secretory, or ciliated) per subject. n = 10 large airway, 7 small airway. *CFTR*$^{+/+}$ and *CFTR* $^{-/-}$ pig samples were grouped together as they had comparable gene expression.

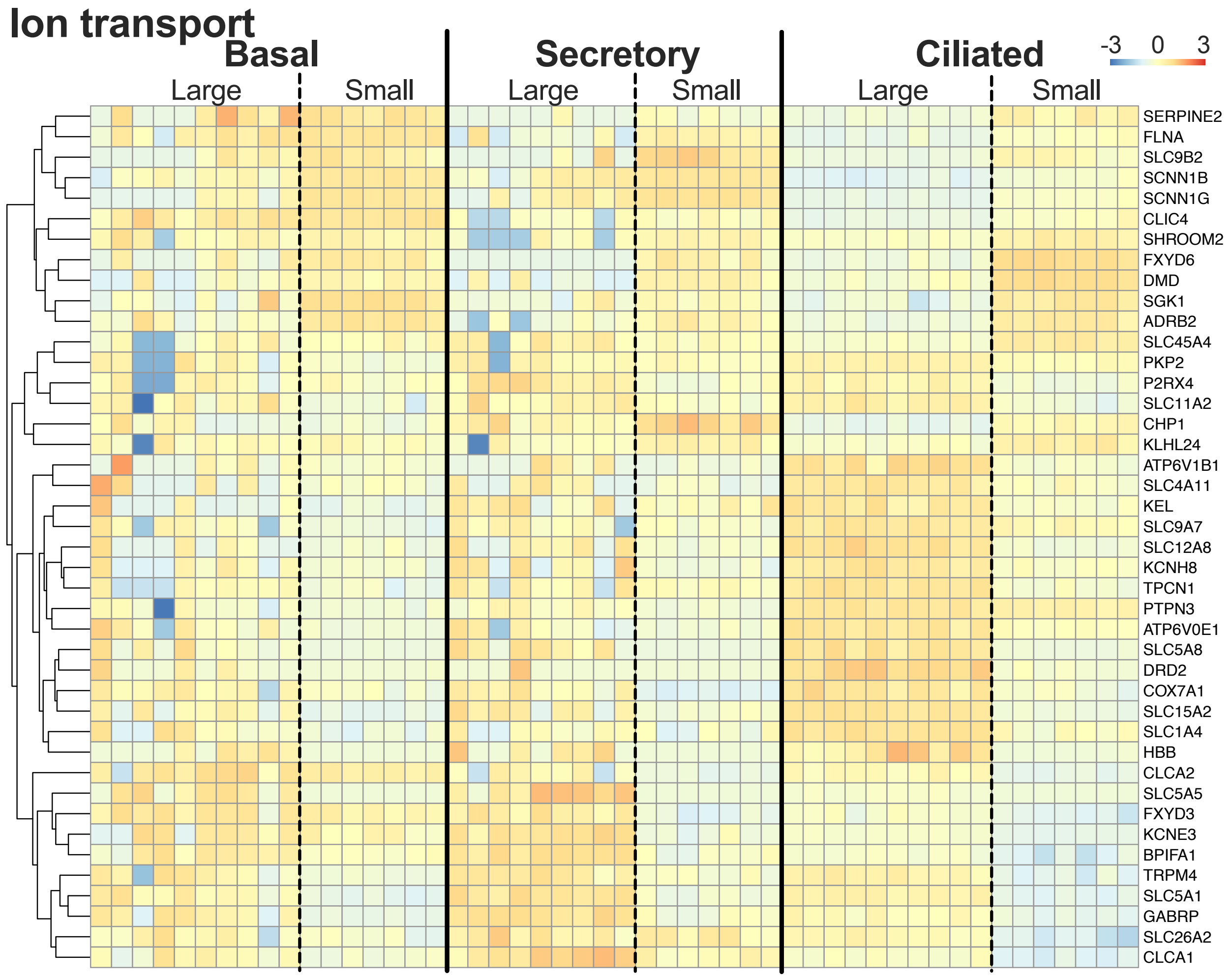


**Supplemental figure 5: Ion transporter and ion transporter regulator genes expression in small and large airway basal, secretory, and ciliated cells at the subject level:** Each column is the normalized/centered average expression per cell type (basal, secretory, or ciliated) per subject. n = 10 large airway, 7 small airway. *CFTR*$^{+/+}$ and *CFTR* $^{-/-}$ pig samples were grouped together as they had comparable gene expression.
